# TSD: Transformers for Seizure Detection

**DOI:** 10.1101/2023.01.24.525308

**Authors:** Yongpei Ma, Chunyu Liu, Maria Sabrina Ma, Yikai Yang, Nhan Duy Truong, Kavitha Kothur, Armin Nikpour, Omid Kavehei

**Affiliations:** School of Biomedical Engineering, The University of Sydney, NSW 2006, Australia; Kids Neuroscience Centre, The Children’s Hospital at Westmead, NSW 2145, Australia; Faculty of Medicine and Health, The University of Sydney, NSW 2006, Australia; Comprehensive Epilepsy Services, Department of Neurology, The Royal Prince Alfred Hospital, Camperdown, NSW 2050, Australia; Faculty of Medicine and Health, The University of Sydney, Sydney, NSW 2006, Australia

**Keywords:** AI, Transformer, Seizure detection, Epilepsy

## Abstract

Epilepsy is a common neurological disorder that substantially deteriorates patients’ safety and quality of life. Electroencephalogram (EEG) has been the golden-standard technique for diagnosing this brain disorder and has played an essential role in epilepsy monitoring and disease management. It is extremely laborious and challenging, if not practical, for physicians and expert humans to annotate all recorded signals, particularly in long-term monitoring. The annotation process often involves identifying signal segments with suspected epileptic seizure features or other abnormalities and/or known healthy features. Therefore, automated epilepsy detection becomes a key clinical need because it can greatly improve clinical practice’s efficiency and free up human expert time to attend to other important tasks. Current automated seizure detection algorithms generally face two challenges: (1) models trained for specific patients, but such models are patient-specific, hence fail to generalize to other patients and real-world situations; (2) seizure detection models trained on large EEG datasets have low sensitivity and/or high false positive rates, often with an area under the receiver operating characteristic (AUROC) that is not high enough for potential clinical applicability.

This paper proposes Transformers for Seizure Detection, which we refer to as TSD in this manuscript. A Transformer is a deep learning architecture based on an encoder-decoder structure and on attention mechanisms, which we apply to recorded brain signals. The AUROC of our proposed model has achieved 92.1%, tested with Temple University’s publically available electroencephalogram (EEG) seizure corpus dataset (TUH). Additionally, we highlight the impact of input domains on the model’s performance. Specifically, TSD performs best in identifying epileptic seizures when the input domain is a time-frequency. Finally, our proposed model for seizure detection in inference-only mode with EEG recordings shows outstanding performance in classifying seizure types and superior model initialization.

## I. Introduction

**E**PILEPSY is a common neurological disorder characterized by recurrent seizures. This neurological disorder has profound social and economic influences on 50 million people worldwide, including stigma, discrimination and the expensive cost of epileptic diagnosis and treatment, with about 70% of these patients lacking proper care [1, 2]. The lack of sophisticated equipment experienced specialists, and unaffordable anti-seizure medicines have led to the “treatment gap”, which means patients with limited access to world-class neurological treatment facilities are not able to receive proper and timely treatments [1]. Therefore, a method with a low cost to detect or predict epileptic seizures is beneficial to improve the life quality of people with epilepsy [3, 4].

Epilepsy severely affects the patient’s life quality. An epileptic seizure is a sudden burst of abnormal electrical activity in the brain that can cause various symptoms. Seizures can vary in severity and duration, and the symptoms a person experiences can depend on the type of seizure. Some common symptoms of seizures include convulsions or muscle spasms, loss of consciousness or awareness, uncontrollable movements of the arms and legs, changes in behaviour or emotion, hallucinations or altered senses, temporary confusion, and more. Although widely unknown, some better-known seizure triggers include sleep deprivation, high fever, stress and certain medications. In people with epilepsy, seizures occur spontaneously and are often recurrent; hence the golden standard in epilepsy diagnosis is to this date around seizure detection [5, 6, 2]. The diagnosis of epilepsy is based on a thorough medical evaluation, which may include a physical exam, neurological exam, and brain imaging tests such as an MRI or CT scan. Electroencephalogram (EEG) remains a core and widely accepted technique in diagnosing and understanding epilepsy which is used to record the brain’s electrical activity and help identify the type of seizures a person is experiencing. The presence of epileptiform abnormalities on an electroencephalogram (EEG) may constitute detection of a seizure [7]. The formal definition of epilepsy, as defined by the International League Against Epilepsy (ILAE) [6], includes the following situations: “(1) At least two unprovoked (or reflex) seizures occurring *>*24h apart; (2) one unprovoked (or reflex) seizure and a probability of further seizures similar to the general recurrence risk (at least 60%) after two unprovoked seizures, occurring over the next ten years.” EEG sometimes is used alongside auxiliary data such as the electrocardiogram (ECG) and audio and video (i.e. video-EEG). It is difficult to determine the exact rate of epilepsy misdiagnosis worldwide, as it can vary depending on various factors, such as the availability of diagnostic resources and the expertise of the healthcare professionals involved. However, misdiagnosis of epilepsy is considered relatively common and significantly consequential for the individual. Literature reported misdiagnosis rates vastly vary with low estimates of 2% and high estimates of over 70%, but a more cited rate is between 20–30% [8]. EEG signal annotation for seizure detection by human experts and specialists is a laborious and time-consuming task; hence, a machine learning tool as an assistant with a human-in-the-loop for final review could achieve outstanding improvements over time and dedicated resources [9].

In this paper, we aim to improve the accuracy of seizure annotation to be used in clinical settings for seizure detection and recognition. The time efficiency of these techniques within an expert-in-the-loop system can be more than ten times relative to manual detection and annotation, as reported in our previous work [9]. Many studies have been conducted by designing an automatic EEG annotation system with machine learning; however, current machine learning algorithms have limitations, including low sensitivity, a high rate of false positives, and strong patient-specificity. We propose a Seizure Detection Transformer (TSD) model. This model was trained and validated on Temple University’s open-source electroencephalogram (EEG) seizure corpus dataset (TUH). TUH is the world’s largest public open-source EEG recordings from a large cohort of people living with epilepsy. On this dataset, we obtained an AUROC of 92.1%, which is 5.4% higher than existing solutions and in this paper, we further elaborate on this result and its significance.

### A. Background

An EEG test is performed by recording electrical activities generated by a high population of neurons (usually) noninvasively and at the surface of the human head by attaching multiple electrodes to the patient’s scalp [10]. EEG comes in many non-invasive and invasive forms and is also a common auxiliary means for experts to localise the epileptic foci (point of origin for focal seizures) and identify the categories of epilepsy, including focal, generalized and unknown [6, 2, 5]. Fisher *et al*. in 2017 revised the classification defined in 1981, which categorized the focal onset into aware and impaired-awareness seizures, the second level of focal onset, and the third level of generalized and unknown onset into the motor and non-motor seizures [5]. Among motor, seizures are automatisms, atonics, clonics, spasms of epilepsy, hyperkinetics, myoclonics, and tonics. Non-motor seizures are also subdivided into autonomic, behaviour arrest, cognitive, emotional, and sensory seizures. In Table II, we summarized seizure types used in this work, along with their profiles and corresponding labels in the dataset.

Clinicians commonly combine long-term EEG monitoring with clinical features of each seizure type to classify onsets and may eventually provide treatment options. For example, focal epilepsy has EEG with focal evolving rhythmic discharges and experiences in the simultaneous or sequential onset of one or more motor or non-motor symptoms [11, 12]. In contrast, the EEG during generalized seizures is bilateral, synchronous, symmetric, and generalized spike-wave complex, and its corresponding clinical characteristic is circadian variations [13]. On the one hand, developing a skilled specialist for EEG reading tends to take several years of practical training, exacerbating the challenge of treatment costs in epilepsy and other neurological disorders. On average, an experienced neurologist, skilled EEG technician, or nurse spends 90-120 minutes carefully reviewing a session of EEG recording, which is usually a 12-hour recording [9]. Deep learning solutions could provide profound benefits on this challenge in an expertin-the-loop style use.

Many recent studies have focused on EEG monitoring using deep learning algorithms. They can be divided into seven types of architecture: (1) convolutional neural networks (CNNs), (2) recurrent neural networks (RNNs), (3) deep belief networks (DBNs), (4) autoencoders (AEs), (5) a new architecture formed by combining CNN with the DBNs or AEs, (6) transformer-based networks; among which 2D-CNNs are the most popular neural network architecture for automated seizure detection [14]. CNN was applied to EEG monitoring and seizure detection diagnosis by transforming EEG signals into one-dimensional or two-dimensional forms and feeding the transformed signals to the CNN model [15, 16, 17, 18, 19, 20, 21]. RNN and its extended models, long short-term memory (LSTM) and gated recurrent units (GRU), were used when the signal has a variation in lengths [22, 23, 24, 25, 26]. AE is an unsupervised deep learning algorithm that combines encoding and decoding blocks to extract features from signals [27, 28, 29, 30]. DBN is also an unsupervised deep learning algorithm that can be considered a generative graphical model with multiple hidden units to reconstruct inputs probabilistically [31, 32]. CNN-RNNs and CNN-AE architectures were similar because they both coupled CNN with RNN or AE modules to diagnose seizures [9, 33, 34, 35, 36, 37, 17, 38]. Transformer-based networks usually add a Transformer module after the CNN convolution module to improve the model’s accuracy [39, 40, 41].

Conventional seizure detection models are often patient-specific and hence extremely difficult, if not impractical, to be generalized [22, 9]. The performances of systems trained on large datasets are often limited due to low area under the receiver operating characteristic curve (AUROC) and high rates of false positives with an acceptable sensitivity to clinicians [9, 42]. It is worth noting that the application of the tool that we are developing plays a significant role in how we will justify the right balance between false alarms and sensitivity. We believe clinical applications that are not in need of a real-time annotation and involve expert reviews based on human psychology and perception tolerate a higher rate of false alarms if it significantly helps sensitivity. We compare our previous study with the state-of-the-art tools in the market in [9].

### B. Novelty

The goal of this work is to overcome the challenge described above. We designed a TSD system to identify epilepsy seizures using pre-recorded EEG signals by short-time Fourier transform (STFT) on the most extensive publicly available EEG dataset, TUH. This paper is part of a newly formed set of papers analysing the use of Transformers on EEG signals and seizure detection to locate and detect epilepsy seizures [43, 41].

Unfortunately, separate Transformer architectures are rarely adopted for signal processing due to the lack of inductive biases, which contributes to the learning process [44, 45]. Thus, current research combined Transformers with convolutional architectures to promote EEG-based diagnoses of seizures [40, 39, 38, 46]. These studies used CNNs to generalize observations and engaged transformers to avoid struggling with modelling context information. However, the stochasticity and complexity of operating environments limit a precise characterization of the inductive bias [47]. This limitation leads to less usefulness of an inductive bias than we imagined. Therefore, the negative impact of this issue can be counterbalanced with a dataset with a large number of data [44].

And compared to other transformer-based networks, our TSD system has the following advantages: (1) the AUROC of our proposed seizure detection model is 92.1%, which is approximately 5% higher than other models that are trained on this dataset, (2) the number of model parameters is small, (3) simple and effective structure, our model combines the transformer with the visual features of EEG, not just processing EEG signals like processing text.

## II. Prerequisite

### A. Montage

The EEG derivatives or channels are arranged logically to form a montage that provides physicians with lateralized and localised information by displaying activity across the whole head [48]. The typical routine EEG recordings are bipolar montages (BM) and referential montages. Our study adopts 17-channel bipolar longitudinal montages with conventional 10-20 placements. The channels are considered between two adjacent electrodes longitudinally between 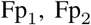, F_3_, F_4_, F_7_, F_8_, C_3_, C_4_, C_*z*_, T_3_, T_4_, P_3_, P_4_, O_1_, O_2_, T_5_, T_6_, and P_*z*_ and F_*z*_ as the reference electrodes.

### B. Transformer

The transformer is a deep learning architecture that uses the multi-head self-attention mechanism to increase the training speed. This technique is commonly applied to parallelized computation.

A transformer model consists of stacked encoders and decoders. An encoder includes a multi-head self-attention module and a position-based fully connected feed-forward network which are connected residually and then their outputs are normalized [49]. Self-attention is shown in this formula:

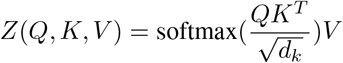

where *Q, K, V* are produced by multiplying the input vectors by three weight matrices *W*_1_, *W*_2_ and *W*_3_, and the *d*_*k*_ means the dimension of the *k*^th^ vector.

Vaswani et al. (2017) proposed the multi-head attention mechanism, which refines the self-attention mechanism. This technique extends the model’s ability to focus on different positions and generates multiple “representation subspaces”. Hence these improvements enhance the performance of the self-attention layer. The multi-head self-attention mechanism employs multiple groups of *Q/K/V* to produce different weight matrices *Z*, which are concatenated as the output of the self-attention layer [49]. Then the model feeds this output into the feed-forward neural network layer. Lastly, the shape of the output matrix is adjusted by multiplying it with an additional weight matrix. The pruned matrix is the input of the feed-forward neural network.

The entire encoding part is formed by stacking multiple encoders. Similarly, the same structure is used in the decoder, which calculates the self-attention score for the output and feeds the output to the forward network. The main difference between an encoder and a decoder is that the decoder consists of a sequence mask to obscure information for future moments.

The final layer of the Transformer model is a fully connected neural network layer and a softmax layer. The linear layer projects the vectors generated by the decoders onto a higher-dimensional vector (logits), where each dimension corresponds to a unique word score. A subsequent softmax layer can compute probabilities in terms of these scores showing in the next equation [50]:

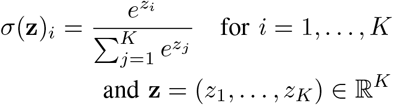

where **z** is input vectors and *K* is its dimension. The word with the highest probability in this dimension is the final output of this time step.

In addition, the Transformer adds a vector with sequential features to each word vector in the input called position vector to save the position information, which is represented by the following formula [49]:

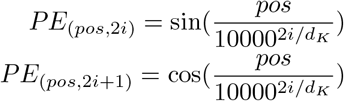

### C. Vision Transformer

Vision Transformers (ViT) applies transformers to visual tasks with a simple effective and strongly scalable model structure. A previous study Dosovitskiy et al. (2020) proposed that ViT with a small size of data usually performs worse than ResNets due to lacking inductive bias. However, it also reported that this could be offset by the increasing training data to improve the performance of ViT which surpasses that of CNN since ViT can obtain better transfer effect in downstream tasks.

## III. Methods

This paper uses the electrode locations and names assigned by the International 10-20 System. When reading an EEG display, we use a representation of a bipolar montage of EEG channels. The next step is to preprocess the input data EEG signals and remove the DC component by the STFT. After that, TSD is established for EEG seizure detection based on the core idea of ViT.

### A. Data Preprocessing

In this paper, we processed EEG signals by the short-time Fourier transform (STFT). Traditionally, the Fourier transform is the most important method for analyzing and processing stationary signals. The temporal domain and frequency domain are two ways to observe a signal. The Fourier transform and its inverse transform convert the signals between the temporal domain and frequency domain [51]. The basic Fourier transform expression is [52]:

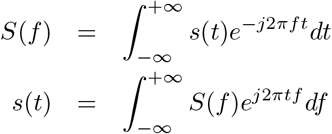

It can be seen that the Fourier transform decomposes the signal into different components as a whole and lacks local information. The Fourier transform cannot combine temporal domain and frequency domain information, which plays an important role in processing non-stationary signals [51].

The short-time Fourier transform (STFT) was proposed to solve this issue. STFT is a widespread method to deal with non-stationary signals, which divides the signal into many small-time intervals (windows), and applies Fourier transform to each one to extract the corresponding frequency [53]. The concatenation of these processed intervals represents the overall temporal spectrum [53]. According to the basic idea, it can be concluded that STFT is designed intuitively for analyzing various processes with approximately the same feature scale rather than multiscale signalling and mutational processes due to the fixed time-frequency window size of STFT [54] Therefore, STFT is suitable for processing raw EEG signals. Here are the expressions of STFT and inverse STFT [53]:

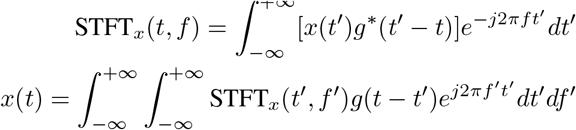

where *x*(*k*) is the original signal, *t* and *g* represent time shift (overlapping part) and window size respectively, and the ∗ represents a complex conjugate. We discretize STFT_*x*_(*t, f*) calculated by continuous STFT in order to achieve it by computer. This equation shows how to obtain the converted signal *F* (*t, f*) [55]:

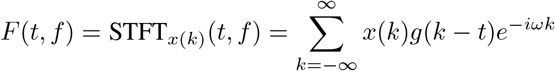

We also can reconstruct the original temporal spectrum by the inverse STFT whose formula is as follows:

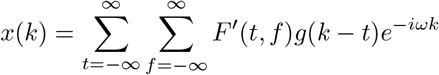

### B. Model structure

This paper uses ViT as the baseline model and adopts the TUH dataset with long-term EEG signals of seizures to train the model to achieve optimal results. We improved the baseline model to apply it for signal processing whose idea is to apply <u>T</u>ransforms for <u>S</u>eizure <u>D</u>etection (TSD).

The architecture of TSD is shown in Fig 1. This model divides the input signal into 200 patches, each with a size of (50, 7). The next step is to project each patch into a fixed-length vector and enter the patch into the Transformer. The subsequent operation of encoders keeps the same as the original Transformer. However, we added a specific token into the input sequence whose corresponding output predicts epileptic seizures. The eventual model is the output module to translate the specific token.

**Fig. 1:**
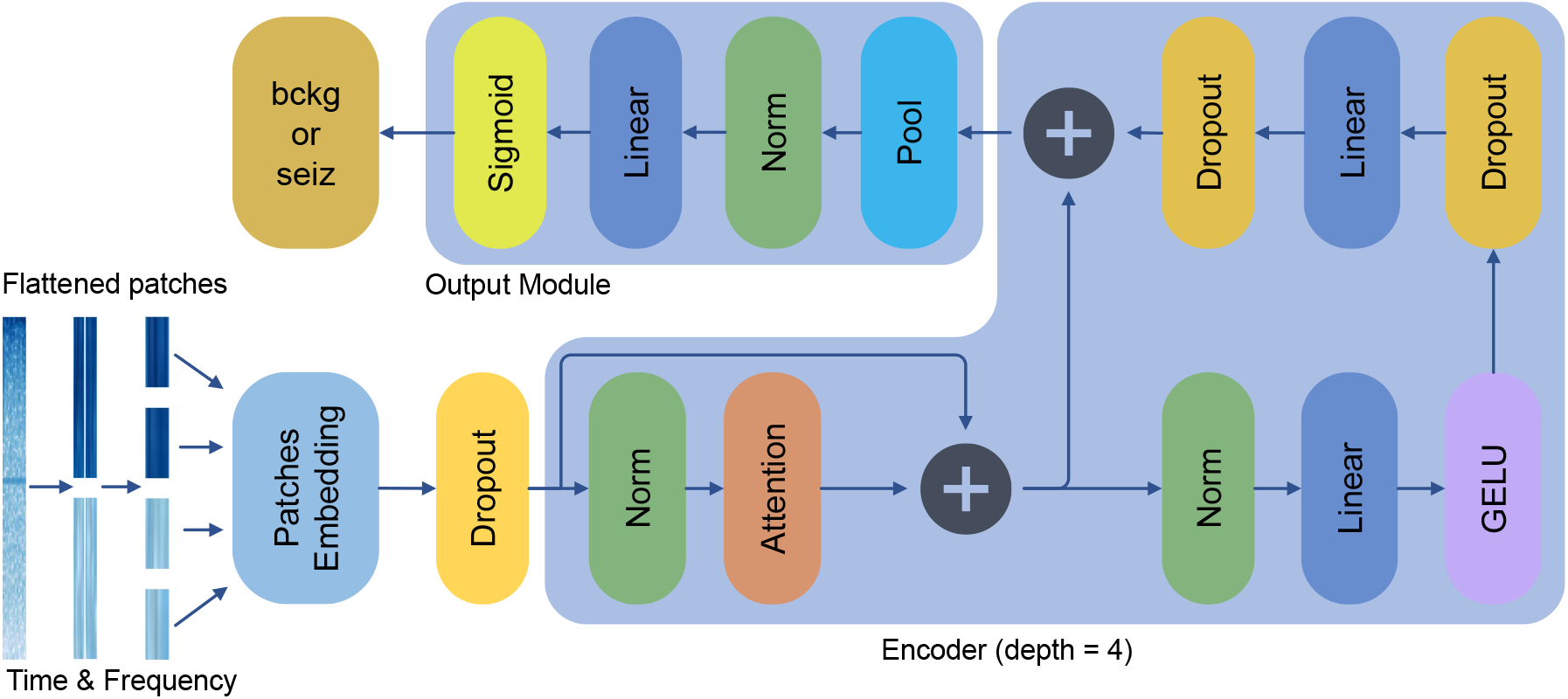
The TSD architecture consists of an input module, encoders, and an output module. The input module converts signals into sequences and embeds them. Encoders employ a multi-head attention mechanism to identify input embeddings. The output module extracts the specific token and obtains predictions.

#### 1) Patch embedding

We divided a signal segment into fixed-size patches, which aims to transform a signal problem into a sequence-to-sequence problem. The input signal size is 5000 × 14, which is split into patches with a resolution of 50 × 7. Thus, each signal segment will generate 200 patches as the input sequence length, and we flatted the 2D patches to a 1D vector with 350 dimensions. The constant latent vector size D = 16 is set as the dimension of the linear projection layer, where we mapped the flattened patches to D-dimension space:

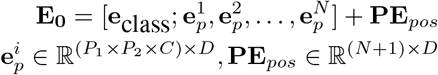

where [] represents the operation of concatenation, *P*_1_ is 50, *P*_2_ is 7, *C* is 1, *D* is 16 and *N* is 200 and the way to add **PE**_*pos*_ is described in the next section. The **e**_class_ is pended to represent the classification **y** of signals.

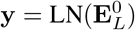

where 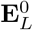 is the output of encoders, and LN is Layer Normalization.

As a result, the dimension of sequential patches is 200 × 16 after passing through the linear projection layer, i.e., there are a total of 200 tokens, and the dimension of each token is 16. In addition, we added a ‘class’ token for outputting the final predicted result. This operation increases the final dimension to 201 × 16.

#### 2) Positional embedding

The positional coding is a standard learnable 1D position embedding that serves as input vector localization records [44]. It can be considered a table with *N* + 1 rows total, each representing a vector with the same dimensions as the input sequence embedding. In this model, we designed a 1-channel and (201, 16) matrix as positional embedding, with internal elements obeying a standard normal distribution.

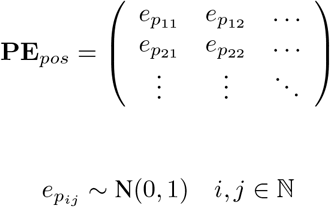

Please note the formula given in the Patch Embedding section, the operation of the positional embedding is a summation rather than a concatenation. Therefore, the dimension of the input embedding sequence remains unchanged, although the position information is added.

#### 3) Encoders

The Encoder is stacked with a couple of single encoder blocks, each of which in this work consists of a multihead self-attention layer (MSA) and a multi-layer perception (MLP). This model applied the multi-head attention mechanism to conduct linear projections to promote the performance of the TSD model [49]. There are 4 heads in this work leading to 4 groups of **Q, K, V** with resolution(201, 4) and concatenated the outputs of these groups.

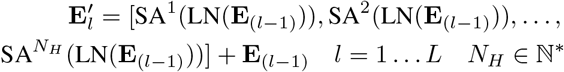

where 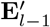 is the output of last block and *N*_*H*_ is the number of heads.

The next part will focus on explaining the structure of the multi-layer perception (MLP). The MLP can be considered as a forward-feed neural network whose learning method is backpropagation [56]. It scales the *x* in terms of proportion. In the proposed model, this block consists of two linear layers (LL) and a non-linear layer with activation function GELU (Gaussian Error Linear Units) [57]:

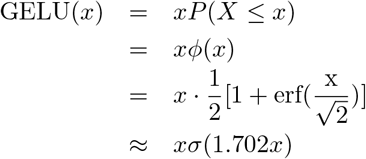

where *x* is the input value of the current neuron and *σ* function is the sigmoid function because of its similarity with the cumulative distribution of the normal distribution. According to Hendrycks and Gimpel (2016), erf() is the Gauss error function, which is defined as:

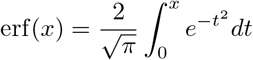

The GELU determines whether x is preserved or not. The results of the multi-head attention layer are normalized by the layer and employed in this module:

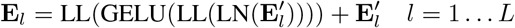

After each part, the present output is layer normalized. At the same time, we added dropout layers to avoid overfitting, whose essence is the achievement of regularization by randomly ignoring half of the neurons.

#### 4) Classification

Furthermore, we did not set decoders after encoders, and the classification results in the output vector will be extracted directly. The output 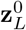 corresponding to the special character ‘class’ will be used as the eventual output of the encoder, representing the signal classification.

### C. Evaluation metrics

In this work, we apply an evaluation metric: Area Under The Curve Receiver Operating Characteristics (AUC-ROC).

#### 1) Area Under The Curve Receiver Operating Characteristics (AUC-ROC)

AUC measures the separability of models and ROC is a probability curve, which reports the ability to the classification of the model [58]. A high AUC means that the model tends to classify correctly. In contrast, a model with an AUC close to 0 shows it reverses the two classes of predictions. An AUC of 0.5 represents the failure of the classification of the model. The AUC-ROC curve shows the change in the ratio of the true-positive rate (TPR) to the false-positive rate (FPR) with the typical and widely known confusion matrix [59].

We take advantage of these indicators to compute TPR and FPR.

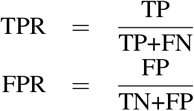

## IV. Experiments

This experiment was conducted on the TUH dataset which was split into a training, a validation and a test set. The data in these subsets were screened for EEG signals from nineteen sensors and the recordings were divided into 12-second segments. Next, these fragments are extracted with time-frequency information using STFT. The model reads the time-frequency graphs in the training set to train and learn the characteristics of EEG signals of epileptic seizures, validates and tests model’s hyperparameters on the validation set, finally evaluates the model predictions on the test set. At the same time, we differentiated the ability of the TSD model to detect various types of seizures. In addition, we illustrated the superiority of the TSD model by comparison of the AUROC between the TSD model and other existing techniques with the same window size and on TUH dataset.

### A. Dataset

We use the Temple University Hospital (TUH) seizure corpus v1.5.4 in our experiments to test the performance of the TSD model. It is the most extensive open-source corpus of the world’s EEG recordings of people with epilepsy. Shah et al. in 2018 described that Temple University Hospital collected, curated and organized clinical EEG data for 14 years. The public can apply this corpus for medical experiments and it can be downloaded from the Neural Engineering Data Consortium. It provides three datasets: a training set, a development set and an evaluation set. In the experiments, we used the training set to train the model. In addition, the dataset mixed by the development set and the evaluation set is divided into two parts, one half is used to verify the efficiency of hyperparameters, and the remaining is used as a test set to test the performance of the model. The detailed subsets of data are summarised in Table I.

**TABLE I:** The summary of TUH dataset

| Dataset attribute | Train | Dev | Eva | Tot |
| --- | --- | --- | --- | --- |
| Files | 4664 | 1832 | 881 | 7377 |
| Sessions | 1174 | 342 | 126 | 1642 |
| Patients | 579 | 53 | 43 | 675 |
| Files with seizures | 872 | 324 | 195 | 1391 |
| Sessions with seizures | 352 | 113 | 63 | 582 |
| Patients with seizures | 208 | 45 | 34 | 287 |
| Number of seizures | 2474 | 1086 | 469 | 4029 |
| Total duration (hours) | 910.3 | 435.5 | 129.9 | 1475.7 |
| absz total duration(hours) | 3.0 | 0.8 | 1.6 | 5.5 |
| cpsz total duration(hours) | 59.4 | 0.2 | 10.0 | 69.6 |
| fnsz total duration(hours) | 188.4 | 71.2 | 31.2 | 290.8 |
| gnsz total duration(hours) | 242.5 | 222.4 | 41.3 | 506.1 |
| mysz total duration(hours) | 7.9 | 0 | 0 | 7.9 |
| spsz total duration(hours) | 8.9 | 0 | 0 | 8.9 |
| tcsz total duration(hours) | 10.6 | 4.5 | 2.6 | 17.7 |
| tnsz total duration(hours) | 2.2 | 0.9 | 0.1 | 3.2 |

In Table I, it can be seen that the proportion of normal EEG signals in the training set is the largest, while the proportion of EEG signals containing epileptic features in the development set and evaluation set is more than that in the training set. This is especially evident in the patient-related data because it is shown that only 36% of patients in the training set were diagnosed with epilepsy, while in the development set and evaluation set, it was as high as 84% and 79% respectively. Such distribution difference indicates that the TSD architecture has strong domain adaptability.

In addition, we conducted data analysis on the TUH dataset in order to explain the feasibility of the TSD model in monitoring EEG signals to detect seizures.

In Fig 2, we counted the proportions of different types of seizure durations in the three data subsets. It can be observed that in each subset, GNSZ has the largest proportion, followed by FNSZ, and other rare epilepsy types have a small proportion. Whereas in the training set and evaluation set, the distribution of GNSZ and FNSZ is similar, accounting for about 47% and 36%, respectively. In the development set, 74% of epilepsy is GNSZ, and only 24% of epilepsy is FNSZ. In addition, we noted that only the training set contains MYSZ and SPSZ, while the development set and evaluation set do not have data on these two types of epilepsy. Therefore, our experiments could not verify the effectiveness of the TSD model on identifying MYSZ and SPSZ.

**Fig. 2:**
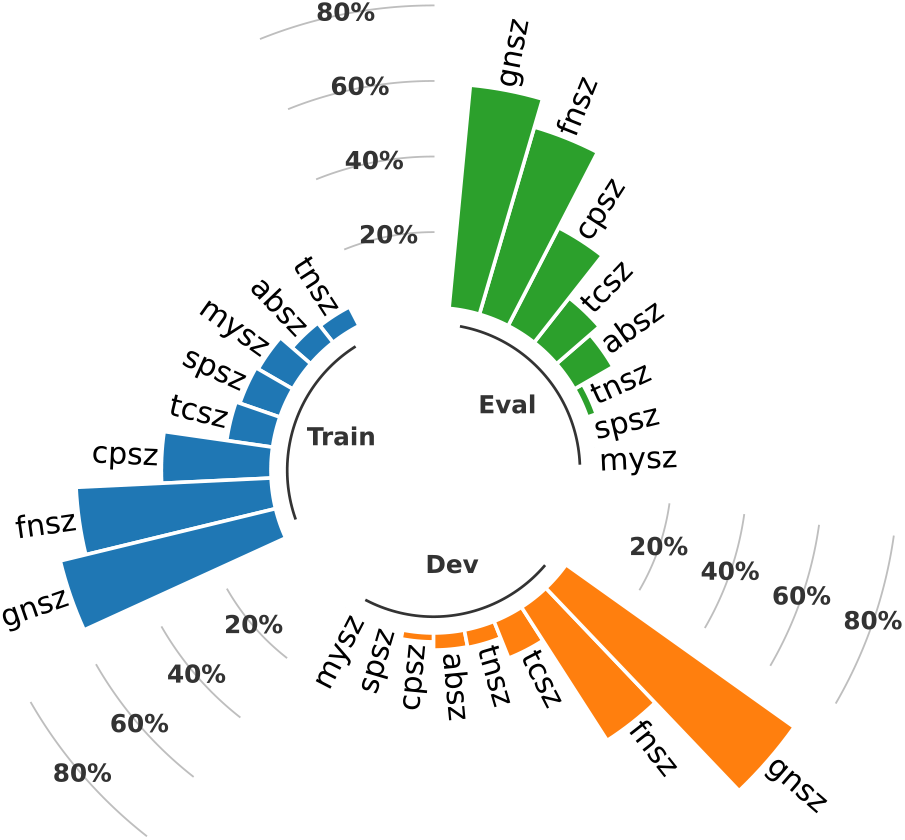
Visualizations of TUH dataset in terms of the distribution of different seizure types in three subsets. Seizure types are explained in Table II. There are in total 7377 files (of 1642 sessions for 675 patients) that we split into training (63%, Train), development (25%, Dev), and evaluation (12%, Eval), of which 18.7% of Train, 17.7% of Dev and 22.1% of Eval are files that contain recorded and documented epileptic seizures.

In Table II, we described the seizure types we classified in this work and which labels were used to annotate their abnormal EEG segments.

**TABLE II:** The summary of classified types of seizures.

| Annotated labels | Seizure type | Description |
| --- | --- | --- |
| **fnsz** | Focal nonspecific seizure. | The origination of Focal seizures is only one hemisphere, and it prefers to propagate the ipsilateral and/or contra-lateral hemisphere [6]. |
| **gnsz** | Generalized seizure. | The origination of Generalized seizure is at a certain point in both hemispheres, and it engages rapidly bilaterally distributed networks [6]. |
| **spsz** | Simple partial seizure. | Simple partial seizure is defined that awareness is preserved during the seizure onsets, and it was revised as aware seizure in 2014 by [6]. |
| **cpsz** | Complex partial seizure. | Complex partial seizure is defined as impaired at any point when the seizure onsets, and it was revised as impaired awareness seizure in 2014 by [6]. |
| **absz** | Absence seizure. | Absence seizure refers to brief, sudden lapses in attention provoked by hyperventilation [6]. |
| **tnsz** | Tonic seizure. | A tonic seizure causes unawareness and muscle contractions of the limbs, which are ongoing from 3 seconds to 2 minutes and a more severe one may include a vibratory component [6]. |
| **tcsz** | Tonic-clonic seizure. | A tonic-clonic seizure occurs in unconscious patients with tonic (increased tone) and a clonic (sustained rhythmic jerking) [6]. |
| **mysz** | Myoclonic seizure. | A myoclonic seizure causes brief muscle contractions lasting more than 30 minutes and with partial awareness [6]. |

### B. Results

We adopted the AUROC as the main evaluation indicator for epileptic detection. The trend of loss and AUROC in the training and validation sets is shown in Fig 3 during the TSD model training with the best results. At the same time, we also conducted other ablation experiments to demonstrate the necessity of various methods in data preprocessing (Appendix A).

**Fig. 3:**
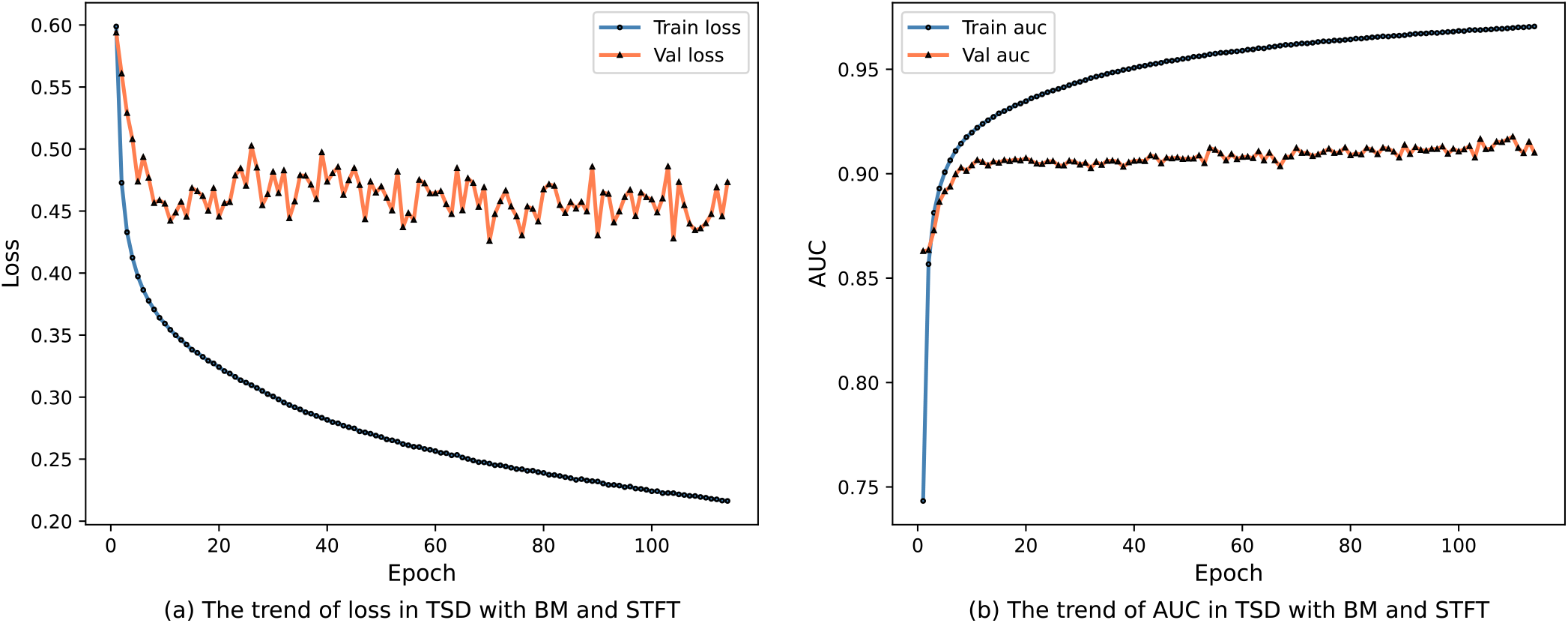
The training process of the proposed model with BM and STFT.

Table III shows the performance of the TSD model and a comparison with the previous studies on the same public dataset with the same window size.

**TABLE III:**
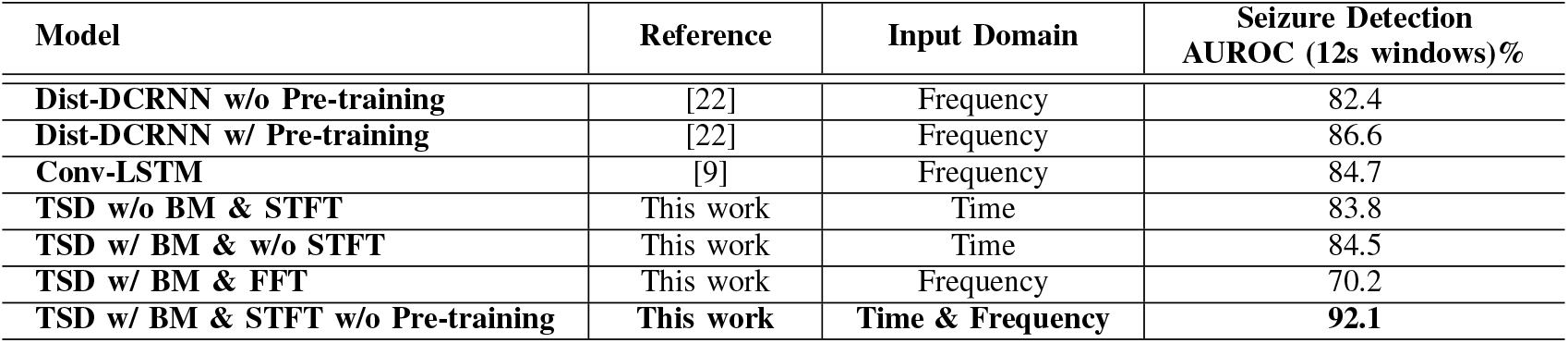
The comparisons of the proposed model with other current models on TUH dataset.

According to Table III, the previous studies performed poorly on large datasets, with a low AUROC, only reached 86.6% at the highest, which is the Dist-DCRNN with Pre-training. Our model achieved an AUCROC of 92.1%. The AUROC of our model for identifying epileptic onset with 12s clip is 5.5% higher than previous state-of-the-art results and we did not pre-train the TSD model. In addition, the improvement in the original model boosted the model performance by 8.3%. We attempted to transfer the input domain to enhance the ability to identify seizures in long-term EEG signals, illustrating that the input domain plays a significant role in seizure detection accuracy. At the same time, the utility of bipolar montages just slightly promotes the AUROC by 0.7%, which is not a obvious performance progress. However, it facilitates the interpretation of the TSD model for seizure detection.

#### 1) Sensitivity to different types of epilepsy

In this section, we examined the sensitivity of the TSD model to different types of epilepsy on the development and evaluation sets to compare the performance of the TSD model for classifying EEG signals. It can be observed in Table IV that for common epilepsy types, the TSD model can detect 96.3% of CPSZ and 91.5% of FNSZ. However, the sensitivity of the GNSZ is only 80.9%. In addition, the model can also sensitively identify rare epilepsy, such as the recall rates of TCSZ and TNSZ are 87.6% and 92.2%, respectively. It is worth noting that although the sensitivity to ABSZ is as high as 100%, considering the short duration of ABSZ in the development and evaluation set, the statistical result about ABSZ has low credibility.

**TABLE IV:** Sensitivity to different types of seizure.

| Seizure type | Sensitivity (%) |
| --- | --- |
| ABSZ | 100 |
| CPSZ | 96.3 |
| FNSZ | 91.5 |
| GNSZ | 80.9 |
| TCSZ | 87.6 |
| TNSZ | 92.2 |

#### 2) Impacts of STFT

In this section, we showed how tuning parameters of STFT affect the accuracy of seizure identification in Fig 4. In the experiments, we refined the window overlap with fixed window size and frequency resolution. In Fig 4, the AUCROC of the proposed TSD model fluctuates with the change of window overlap until reaching the optimal value of 92.1% when the overlap rate is 20%.

**Fig. 4:**
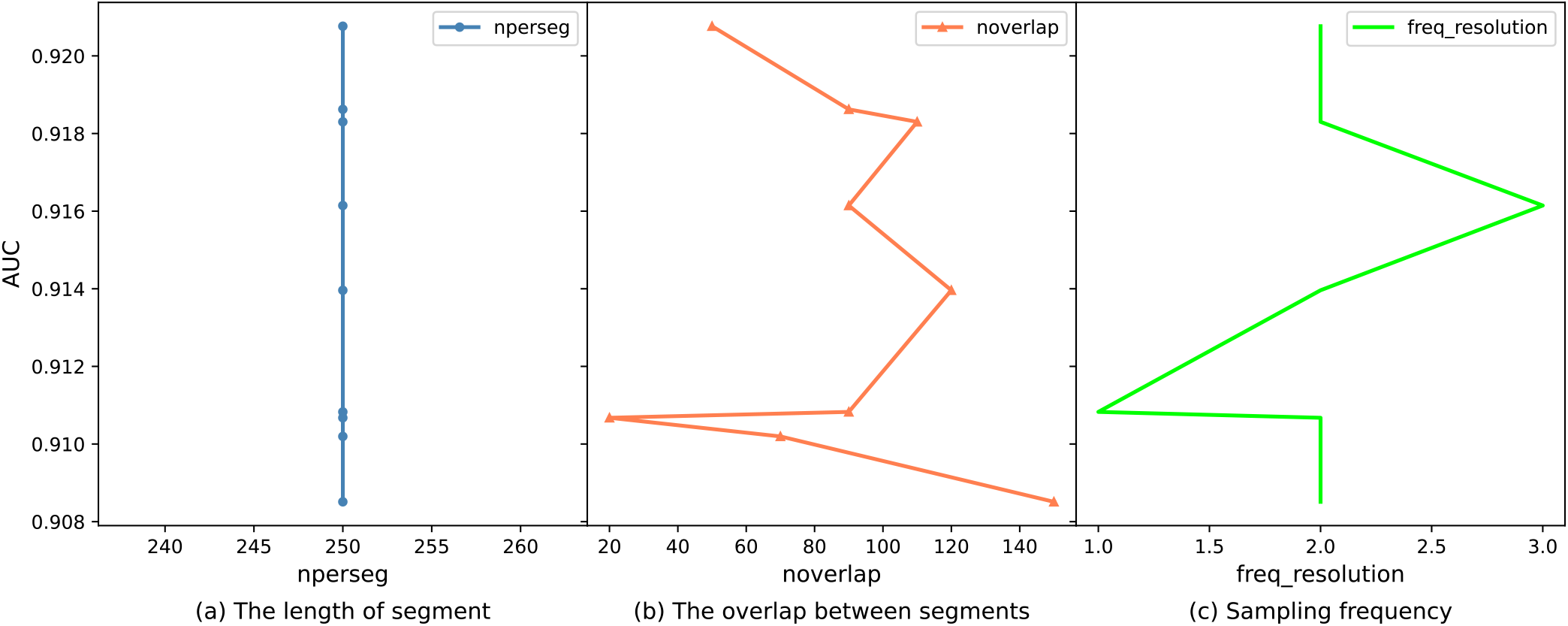
Results with different STFT parameters. Fig (a) shows the changing trend of AUC with the length of a segment. Fig (b) shows the trend of AUC with the increase in overlap rate. Fig (c) documents the AUC of the proposed model with different frequency resolutions.

### 3) Impacts of input domain

We compared the effect of different input domains on seizure detection results. We applied the Fast Fourier Transform (FFT) for the extraction of frequency information. FFT is a common method to process signals. The results show that the extraction of temporal domain information is more critical for the model to learn the EEG features of epileptic patients than the frequency domain information since the change in the input domain of the model from the temporal domain to the frequency domain decreased the AUROC of the model from 84.5% to 70.2%. However, when we simultaneously extracted the time-frequency information of the input EEG signal and fed it as the model input, the model performance for seizure detection increased to 92.1%. Such advances suggest that connection in the time-frequency domain promotes the model’s effectiveness, which is consistent with the conclusion of [22].

#### 4) Superiority

The AUROC of our baseline model without preprocessing the data is only 2.8% lower than the previous state-of-the-art. Notably, Tang et al. in 2021 used a self-supervised pre-training approach to improve model accuracy. At the same time, Tang et al. also used GNN technology to deal with the non-Euclidean structure of nodes to optimize the model performance, which consumes a lot of time and resources. Yang et al. in 2022 post-processed the output results. The output results of the Conv-LSTM algorithm is sent to a postprocessing algorithm to review the results in real-time and make a decision. Our model improves on the previously published model, indicating the superior model initialization and strong large-scale learning ability of TSD. Therefore, we consider that our model can scale and, upon application to clinics, reduce the economic and human burden of long-term EEG monitoring.

## V. Conclusion

In conclusion, we proposed a TSD algorithm for learning EEG recordings and monitoring epilepsy. We also validated the ability of the TSD model to detect epilepsy on a large corpus of public EEG recordings TUH. We significantly improved the performance of state-of-the-art AI algorithms for seizure detection and classification and compared the impact of the data-preprocessing methods and the input domain on the model’s ability to identify seizures. In addition, we demonstrated that our model has excellent model initialization and is more conducive to overcoming the patient-specific problems of existing seizure detection instruments.

In the future, it is worth using the proposed model for EEG-based epilepsy classification to locate the lesion location of epileptic seizures. This attempt can facilitate clinicians’ current dilemma in the seizure onset area. Furthermore, Our model did not distinguish between patient age groups to verify whether the TSD model has the same effect on different age groups. In fact, real-world seizure detection in neonates and children is a more challenging limitation. Therefore, our future direction will focus on improving the model for application in neonatal and childhood epilepsy screening.

## Appendix A Setup in ablation experiments

We conducted three ablation experiments on TSD with different data-processing methods. (1) We split the raw EEG signals in the dataset into 12s clips. Then these segments are directly used as the input of the TSD model. (2) We built a bipolar montage with the original signal after we segmented the original signal. (3) On the basis of bipolar montage, we used FFT to convert the information in each 12s segment from the time domain to the frequency domain. We demonstrated the trend of AUC during training for these three ablation experiments in Fig 5.

**Fig. 5:**
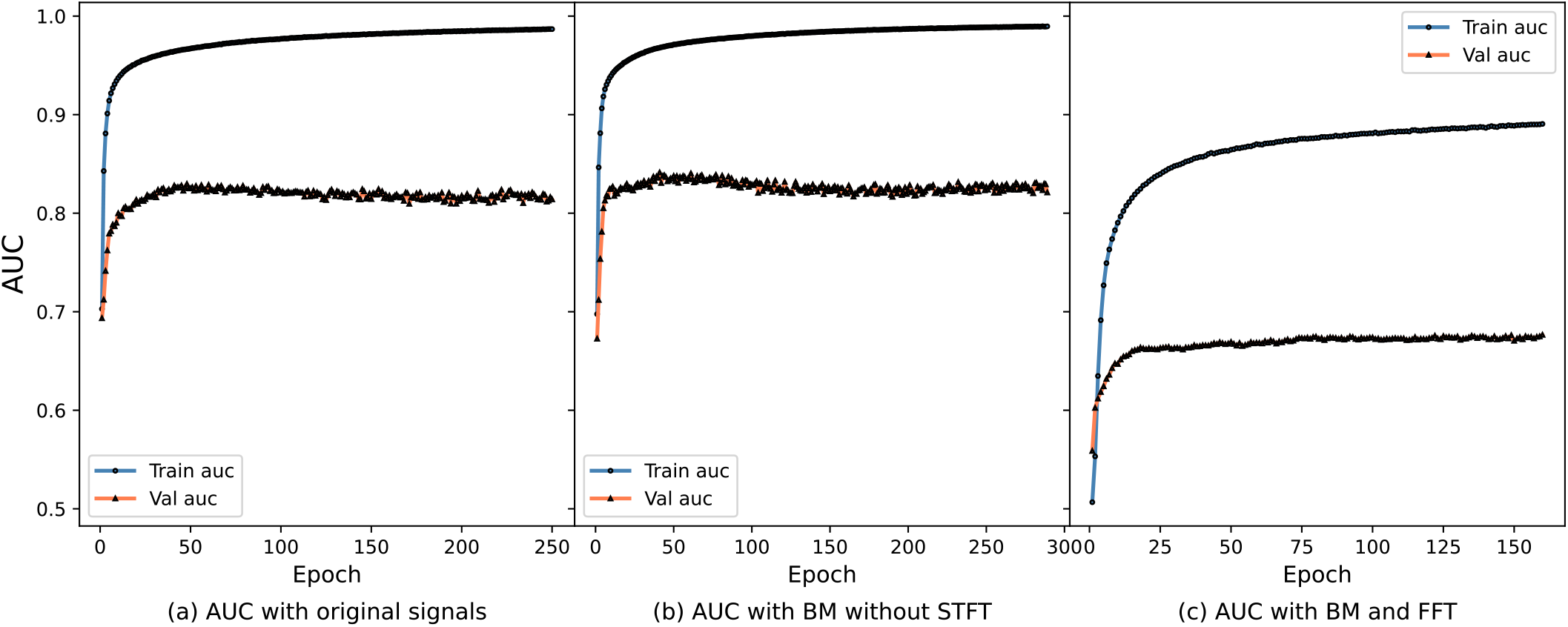
The training process of the proposed model in all ablation experiments

## Notes

### Competing Interest Statement

The authors have declared no competing interest.

### Summary of Updates

Add two authors

## References

[1] Abdul Wahab. Difficulties in treatment and management of epilepsy and challenges in new drug development. Pharmaceuticals, 3(7):2090–2110, 2010.

[2] Nhan Duy Truong. Epileptic Seizure Detection and Forecasting Ecosystems. PhD Dissertation, The University of Sydney, 2020.

[3] Epilepsy Foundation. Epilepsy Innovation Institute (EI2) Community Survey, 2016.

[4] Sonya B Dumanis, Jaqueline A French, Christophe Bernard, Gregory A Worrell, and Brandy E Fureman. Seizure forecasting from idea to reality. Outcomes of the my seizure gauge epilepsy innovation institute workshop. eNeuro, 4(6), 2017.

[5] Robert S Fisher, J Helen Cross, Carol D’souza, Jacqueline A French, Sheryl R Haut, Norimichi Higurashi, Edouard Hirsch, Floor E Jansen, Lieven Lagae, Solomon L Moshé, et al. Instruction manual for the ILAE 2017 operational classification of seizure types. Epilepsia, 58(4):531–542, 2017.

[6] Robert S Fisher, Carlos Acevedo, Alexis Arzimanoglou, Alicia Bogacz, J Helen Cross, Christian E Elger, Jerome Engel Jr, Lars Forsgren, Jacqueline A French, Mike Glynn, et al. ILAE official report: a practical clinical definition of epilepsy. Epilepsia, 55(4):475–482, 2014.

[7] Samuel L Bridgers, Peter B Wade, and John S Ebersole. Estimating the importance of epileptiform abnormalities discovered on cassette electroencephalographic monitoring. Archives of neurology, 46(10):1077–1079, 1989.

[8] Maria Meritxell Oto. The misdiagnosis of epilepsy: appraising risks and managing uncertainty. Seizure, 44: 143–146, 2017.

[9] Yikai Yang, Nhan Duy Truong, Christina Maher, Armin Nikpour, and Omid Kavehei. Continental generalization of a human-in-the-loop AI system for clinical seizure recognition. Expert Systems with Applications, 207: 118083, 2022.

[10] Ernst Niedermeyer and FH Lopes da Silva. Electroencephalography: basic principles, clinical applications, and related fields. Lippincott Williams & Wilkins, 2005.

[11] Jeffrey W. Britton, Lauren C. Frey, Jennifer L. Hopp, Pearce Korb, Mohamad Z. Koubeissi, William E. Lievens, Elia M. Pestana-Knight, and Erik K. St. Louis. Electroencephalography (EEG): An introductory text and atlas of normal and abnormal findings in adults, children, and infants, 2016. URL http://europepmc.org/books/NBK390354.

[12] Ronit M Pressler, Maria Roberta Cilio, Eli M Mizrahi, Solomon L Moshé, Magda L Nunes, Perrine Plouin, Sampsa Vanhatalo, Elissa Yozawitz, Linda S de Vries, Kollencheri Puthenveettil Vinayan, et al. The ILAE classification of seizures and the epilepsies: Modification for seizures in the neonate. position paper by the ILAE task force on neonatal seizures. Epilepsia, 62(3):615–628, 2021.

[13] Udaya Seneviratne, Mark J Cook, and Wendyl Jude D’Souza. Electroencephalography in the diagnosis of [1] genetic generalized epilepsy syndromes. Frontiers in neurology, 8:499, 2017.

[14] Afshin Shoeibi, Marjane Khodatars, Navid Ghassemi, Mahboobeh Jafari, Parisa Moridian, Roohallah Alizadehsani, Maryam Panahiazar, Fahime Khozeimeh, Assef Zare, Hossein Hosseini-Nejad, et al. Epileptic seizures detection using deep learning techniques: a review. International Journal of Environmental Research and Public Health, 18(11):5780, 2021.

[15] Khaled Saab, Jared Dunnmon, Christopher Ré, Daniel Rubin, and Christopher Lee-Messer. Weak supervision as an efficient approach for automated seizure detection in electroencephalography. NPJ digital medicine, 3(1): 1–12, 2020.

[16] Mustafa Talha Avcu, Zhuo Zhang, and Derrick Wei Shih Chan. Seizure detection using least EEG channels by deep convolutional neural network. In ICASSP 2019-2019 IEEE international conference on acoustics, speech and signal processing (ICASSP), pages 1120–1124. IEEE, 2019.

[17] Ahmed M Abdelhameed, Hisham G Daoud, and Magdy Bayoumi. Epileptic seizure detection using deep convolutional autoencoder. In 2018 IEEE International Workshop on Signal Processing Systems (SiPS), pages 223–228. IEEE, 2018.

[18] M Shamim Hossain, Syed Umar Amin, Mansour Alsulaiman, and Ghulam Muhammad. Applying deep learning for epilepsy seizure detection and brain mapping visualization. ACM Transactions on Multimedia Computing, Communications, and Applications (TOMM), 15(1s): 1–17, 2019.

[19] Rui Zuo, Jing Wei, Xiaonan Li, Chunlin Li, Cui Zhao, Zhaohui Ren, Ying Liang, Xinling Geng, Chenxi Jiang, Xiaofeng Yang, et al. Automated detection of highfrequency oscillations in epilepsy based on a convolutional neural network. Frontiers in computational neuroscience, 13:6, 2019.

[20] Ian C Covert, Balu Krishnan, Imad Najm, Jiening Zhan, Matthew Shore, John Hixson, and Ming Jack Po. Temporal graph convolutional networks for automatic seizure detection. In Machine Learning for Healthcare Conference, pages 160–180. PMLR, 2019.

[21] Bassem Bouaziz, Lotfi Chaari, Hadj Batatia, and Antonio Quintero-Rincón. Epileptic seizure detection using a convolutional neural network. In Digital Health Approach for Predictive, Preventive, Personalised and Participatory Medicine, pages 79–86. Springer, 2019.

[22] Siyi Tang, Jared Dunnmon, Khaled Kamal Saab, Xuan Zhang, Qianying Huang, Florian Dubost, Daniel Rubin, and Christopher Lee-Messer. Self-supervised graph neural networks for improved electroencephalographic seizure analysis. In International Conference on Learning Representations, 2021.

[23] Xuhui Chen, Jinlong Ji, Tianxi Ji, and Pan Li. Cost-sensitive deep active learning for epileptic seizure detection. In Proceedings of the 2018 ACM International Conference on Bioinformatics, Computational Biology, and Health Informatics, pages 226–235, 2018.

[24] Kosuke Fukumori, Hoang Thien Thu Nguyen, Noboru Yoshida, and Toshihisa Tanaka. Fully data-driven convolutional filters with deep learning models for epileptic spike detection. In ICASSP 2019-2019 IEEE international conference on acoustics, speech and signal processing (ICASSP), pages 2772–2776. IEEE, 2019.

[25] Lasitha Vidyaratne, Alexander Glandon, Mahbubul Alam, and Khan M Iftekharuddin. Deep recurrent neural network for seizure detection. In 2016 International Joint Conference on Neural Networks (IJCNN), pages 1202–1207. IEEE, 2016.

[26] Minxing Geng, Weidong Zhou, Guoyang Liu, Chaosong Li, and Yanli Zhang. Epileptic seizure detection based on stockwell transform and bidirectional long short-term memory. IEEE Transactions on Neural Systems and Rehabilitation Engineering, 28(3):573–580, 2020.

[27] Meysam Golmohammadi, Amir Hossein Harati Nejad Torbati, Silvia Lopez de Diego, Iyad Obeid, and Joseph Picone. Automatic analysis of EEGs using big data and hybrid deep learning architectures. Frontiers in human neuroscience, 13:76, 2019.

[28] Ali Emami, Naoto Kunii, Takeshi Matsuo, Takashi Shinozaki, Kensuke Kawai, and Hirokazu Takahashi. Autoencoding of long-term scalp electroencephalogram to detect epileptic seizure for diagnosis support system. Computers in biology and medicine, 110:227–233, 2019.

[29] Vinit Shah, Meysam Golmohammadi, Saeedeh Ziyabari, Eva Von Weltin, Iyad Obeid, and Joseph Picone. Optimizing channel selection for seizure detection. In IEEE signal processing in medicine and biology symposium (SPMB), pages 1–5. IEEE, 2017.

[30] Ye Yuan, Guangxu Xun, Kebin Jia, and Aidong Zhang. A multi-view deep learning method for epileptic seizure detection using short-time Fourier transform. In Proceedings of the 8th ACM International Conference on Bioinformatics, Computational Biology, and Health Informatics, pages 213–222, 2017.

[31] Thanh Xuyen Le, Trung Thanh Le, Van Viet Dinh, Quoc Long Tran, Linh Trung Nguyen, and Duc Thuan Nguyen. Deep learning for epileptic spike detection. VNU Journal of Science: Computer Science and Communication Engineering, 33(2):1–13, 2018.

[32] JT Turner, Adam Page, Tinoosh Mohsenin, and Tim Oates. Deep belief networks used on high resolution multichannel electroencephalography data for seizure detection. In 2014 aaai spring symposium series, 2014.

[33] Subhrajit Roy, Isabell Kiral-Kornek, and Stefan Harrer. Deep learning enabled automatic abnormal EEG identification. In 2018 40th Annual International Conference of the IEEE Engineering in Medicine and Biology Society (EMBC), pages 2756–2759. IEEE, 2018.

[34] Zhou Fang, Howan Leung, and Chiu Sing Choy. Spatial temporal GRU convnets for vision-based real time epileptic seizure detection. In 2018 IEEE 15th International Symposium on Biomedical Imaging (ISBI 2018), pages 1026–1029. IEEE, 2018.

[35] Gwangho Choi, Chulkyun Park, Junkyung Kim, Kyoungin Cho, Tae-Joon Kim, HwangSik Bae, Kyeongyuk [1] Min, Ki-Young Jung, and Jongwha Chong. A novel multi-scale 3D CNN with deep neural network for epileptic seizure detection. In IEEE International Conference on Consumer Electronics (ICCE), pages 1–2. IEEE, 2019.

[36] Ye Yuan, Guangxu Xun, Kebin Jia, and Aidong Zhang. A multi-view deep learning framework for EEG seizure detection. IEEE journal of biomedical and health informatics, 23(1):83–94, 2018.

[37] Tingxi Wen and Zhongnan Zhang. Deep convolution neural network and autoencoders-based unsupervised feature learning of EEG signals. IEEE Access, 6:25399–25410, 2018.

[38] Siyi Tang, Jared A Dunnmon, Liangqiong Qu, Khaled K Saab, Christopher Lee-Messer, and Daniel L Rubin. Spatiotemporal modeling of multivariate signals with graph neural networks and structured state space models. arXiv preprint arXiv:2211.11176, 2022.

[39] Jonathan Pedoeem, Guy Bar Yosef, Shifra Abittan, and Sam Keene. Tabs: Transformer based seizure detection. In Biomedical Sensing and Analysis, pages 133–160. Springer, 2022.

[40] Yulin Sun, Weipeng Jin, Xiaopeng Si, Xingjian Zhang, Jiale Cao, L. Wang, Shaoya Yin, and Dong Ming. Continuous seizure detection based on Transformer and long-term iEEG. IEEE Journal of Biomedical and Health Informatics, 26(11):5418–5427, 2022. doi: 10.1109/JBHI.2022.3199206.

[41] Wei Yan Peh, Prasanth Thangavel, Yuanyuan Yao, John Thomas, Yee-Leng Tan, and Justin Dauwels. Six-center assessment of CNN-Transformer with belief matching loss for patient-independent seizure detection in EEG. International Journal of Neural Systems, 2022.

[42] Yikai Yang, Nhan Duy Truong, Jason K Eshraghian, Armin Nikpour, and Omid Kavehei. Weak selfsupervised learning for seizure forecasting: a feasibility study. Royal Society Open Science, 9(8):220374, 2022.

[43] Paola Busia, Andrea Cossettini, Thorir Mar Ingolfsson, Simone Benatti, Alessio Burrello, Moritz Scherer, Matteo Antonio Scrugli, Paolo Meloni, and Luca Benini. EEGformer: Transformer-based epilepsy detection on raw EEG traces for low-channel-count wearable continuous monitoring devices. In IEEE Biomedical Circuits and Systems Conference (BioCAS), pages 640–644. IEEE, 2022.

[44] Alexey Dosovitskiy, Lucas Beyer, Alexander Kolesnikov, Dirk Weissenborn, Xiaohua Zhai, Thomas Unterthiner, Mostafa Dehghani, Matthias Minderer, Georg Heigold, × Sylvain Gelly, et al. An image is worth 16 × 16 words: Transformers for image recognition at scale. arXiv preprint arXiv:2010.11929, 2020.

[45] Fozia Mehboob, Abdul Rauf, Richard Jiang, Abdul Khader Jilani Saudagar, Khalid Mahmood Malik, Muhammad Badruddin Khan, Mozaherul Hoque Abdul Hasnat, Abdullah AlTameem, and Mohammed AlKhathami. Towards robust diagnosis of COVID-19 using vision self-attention transformer. Scientific Reports, 12(1):1–12, 2022.

[46] Chang Li, Xiaoyang Huang, Rencheng Song, Ruobing Qian, Xiang Liu, and Xun Chen. EEG-based seizure prediction via Transformer guided CNN. Measurement, 203:111948, 2022.

[47] Eyke Hullermeier, Thomas Fober, and Marco Mernberger. Inductive bias. In Werner Dubitzky, Olaf Wolkenhauer, Kwang-Hyun Cho, and Hiroki Yokota, editors, Encyclopedia of Systems Biology, pages 1018–1019. Springer, New York, 2013. ISBN 978-1-4419-9863-7.

[48] Jayant N Acharya and Vinita J Acharya. Overview of EEG montages and principles of localization. Journal of Clinical Neurophysiology, 36(5):325–329, 2019.

[49] Ashish Vaswani, Noam Shazeer, Niki Parmar, Jakob Uszkoreit, Llion Jones, Aidan N Gomez, Lukasz Kaiser, and Illia Polosukhin. Attention is all you need. Advances in neural information processing systems, 30, 2017.

[50] Ian Goodfellow and Aaron Courvill. Softmax units for multinoulli output distributions, page 180–184. MIT Press, 2016.

[51] Daniel Potts, Gabriele Steidl, and Manfred Tasche. Fast Fourier transforms for nonequispaced data: A tutorial. Modern sampling theory, pages 247–270, 2001.

[52] Jean Baptiste Joseph Fourier, Gaston Darboux, et al. Theórie analytique de la chaleur, volume 504. Didot Paris, 1822.

[53] Daniel Griffin and Jae Lim. Signal estimation from modified short-time Fourier transform. IEEE Transactions on Acoustics, Speech, and Signal Processing, 32(2):236– 243, 1984.

[54] Jung Jun Lee, Sang Min Lee, In Young Kim, Hong Ki Min, and Seung Hong Hong. Comparison between short time fourier and wavelet transform for feature extraction of heart sound. In Multimedia Technology for Asia-Pacific Information Infrastructure (TENCON), volume 2, pages 1547–1550. IEEE, 1999.

[55] Alan V Oppenheim. Discrete-time signal processing. Pearson Education India, 1999.

[56] Simon Haykin and N Network. A comprehensive foundation. Neural networks, 2(2004):41, 2004.

[57] Dan Hendrycks and Kevin Gimpel. Gaussian error linear units (gelus). arXiv preprint arXiv:1606.08415, 2016.

[58] James A Hanley and Barbara J McNeil. The meaning and use of the area under a receiver operating characteristic (ROC) curve. Radiology, 143(1):29–36, 1982.

[59] Stephen V Stehman. Selecting and interpreting measures of thematic classification accuracy. Remote sensing of Environment, 62(1):77–89, 1997.

[60] Vinit Shah, Eva Von Weltin, Silvia Lopez, James Riley McHugh, Lillian Veloso, Meysam Golmohammadi, Iyad Obeid, and Joseph Picone. The temple university hospital seizure detection corpus. Frontiers in neuroinformatics, 12:83, 2018.

